# Tracking movements of an endangered bird using mark-recapture based on DNA-tagging

**DOI:** 10.1101/2023.01.12.523713

**Authors:** María José Bañuelos, María Morán-Luis, Patricia Mirol, Mario Quevedo

## Abstract

Knowing the location and movements of individuals at various temporal and spatial scales is an important facet of behaviour and ecology. In threatened populations, movements that would ensure adequate genetic flow and long term population viability are often challenged by habitat fragmentation. It is also in those endangered populations where capturing and handling individuals to equip them with transmitters or to obtain tissue samples may present additional logistical challenges. DNA-tagging, i.e. individual identification of samples obtained via non-invasive approaches, can reveal certain movement patterns. We used faecal material genetically assigned to individuals to indirectly track movements of a large-bodied, endangered forest bird, Cantabrian capercaillie (*Tetrao urogallus cantabricus*), for three consecutive mating seasons. We identified 127 individuals, and registered movements of 70 of them (22 females, 48 males). Most movements were relatively short for capercaillie, mostly concentrated around display areas. We did not find differences in movement distances between females and males within mating seasons, or between them. Several longer, inter-valley movements up to 9.9 km of planimetric distance linked distant display areas, showing that both females and males of Cantabrian capercaillie were able to move through the landscape, complementing previous studies on gene flow. Those longer movements may be taking birds outside of the study area, and into historical capercaillie territories, which still include substantial forest cover. Tracking animals via DNA tagging, particularly those on endangered populations, showed clear advantages like non-intrusiveness and potential for sample sizes much larger than via direct handling. However, it also misses out on direct observation and natural history, which would provide key information like social status and timing of movements.

## Introduction

The study of movement patterns at various temporal and spatial scales provides basis for understanding behaviour, ecology and, ultimately, aspects of the conservation status of most animals (Börger, Dalziel, & Fryxell, 2008). Some animal movements are tightly associated to their specific habitat requirements, and in those cases navigating unfamiliar and unsuitable habitats might increase risk exposure, compromising survival (Yoder et al. 2004; Bonte et al. 2012). As consequence, the loss and fragmentation of habitat that affects most terrestrial ecosystems (e.g. Watson et al. 2018) may result in appearance of dispersal barriers, which affect population dynamics and gene flow via reduced connectivity (Ricketts 2001; Caplat et al. 2016). Indeed, habitat loss and fragmentation of previous distribution ranges may yield smaller, subdivided populations, if the loss of connectivity results in less frequent movements, and limits effective dispersal. Those subdivided populations are more vulnerable to stochastic events that could restrict their probability of survival (Lens et al., 2002).

The combination of the species’ movement ability and the presence and shape of remnant habitat fragments together with the characteristics of the surrounding matrix, determine the species’ responses to landscape changes. Thus fine-scale movement behaviour of species of conservation interest should be incorporated in conservation planning and management, particularly when habitat restoration is feasible (Lechner et al., 2015). Yet, such dataset are not easy to obtain, and tend to be replaced by assumptions and simplifications (e.g. Southwell et al. 2008) derived from data from other populations, or similar species. However, movements are often partially determined by landscape characteristics, and by distinct selective pressures on different populations (Baguette et al. 2013), so it is not always safe assuming that populations of a given species would show equivalent movement patterns throughout their distribution range. For instance, the established knowledge on a species’ behaviour and habitat requirements may have been obtained from parts of its distribution range less affected by habitat loss and fragmentation. Yet, in fragmented and peripheral areas of a distribution range, the quality of what may appear as secondary habitat is possibly key for the species’ survival (Channel and Lomolino 2000; Blanco-Fontao et al. 2010). This is particularly general for those areas within a species’ range where dispersal movements depend on fine-scale structural elements, acting as stepping stones (Lechner et al. 2015).

We studied movements of an endangered forest bird, Cantabrian capercaillie *Tetrao urogallus cantabricus*, living at the southern edge of the species range, under the influence of Atlantic climate (Olson et al. 2001; Cervellini et al. 2020). This capercaillie population declined severely from its known historical range in the last third of the 20th century (Pollo et al. 2005; Storch et al. 2006), and its viability appears compromised (Bañuelos et al. 2019). It has been recently listed as critically endangered in Spain (Ministerio para la Transición Ecológica 2018). Therefore, to study them, it was advisable to use minimally intrusive techniques, like tagging of individuals based on their DNA, left behind in shed tissues or scats. DNA tagging, where the repeated occurrence of a genotype is a direct indicator of movement, has been used to study movements at various spatial and temporal scales (Palsbøll et al. 1997; Palsbøll 1999). It allows monitoring a wider representation of the variability of individuals in a population, albeit with with fewer observations per individual than direct tracking techniques.

The remnant distribution range of Cantabrian capercaillie includes some relatively large patches of forest and shrubland above the treeline, the latter being especially relevant for females with broods (Bañuelos et al. 2008). Yet, these habitats are embedded in a patchwork of disturbed landscape, including former forest clearings and burns in various states of secondary succession, pastures, roads, villages, and industrial activities like mountaintop mining; several known display areas are less than 500 meters from the edge of those mining operations. We wanted to know how capercaillie moved in such landscape, and whether they would show regular, relatively long movements.

## Methods

### Field survey

The study area (Fig. 1) lays within the region of broadleaf forests under Atlantic influence (e.g. Olson et al. 2001). The area is rugged, with elevation ranging from 450 to 2000 m a.s.l. Larger patches of relatively old forest are restricted to the higher slopes of the mountain range, mirroring a widespread pattern of landscape configuration related to human impacts (Sandel and Svenning 2013).

**Figure 1.**
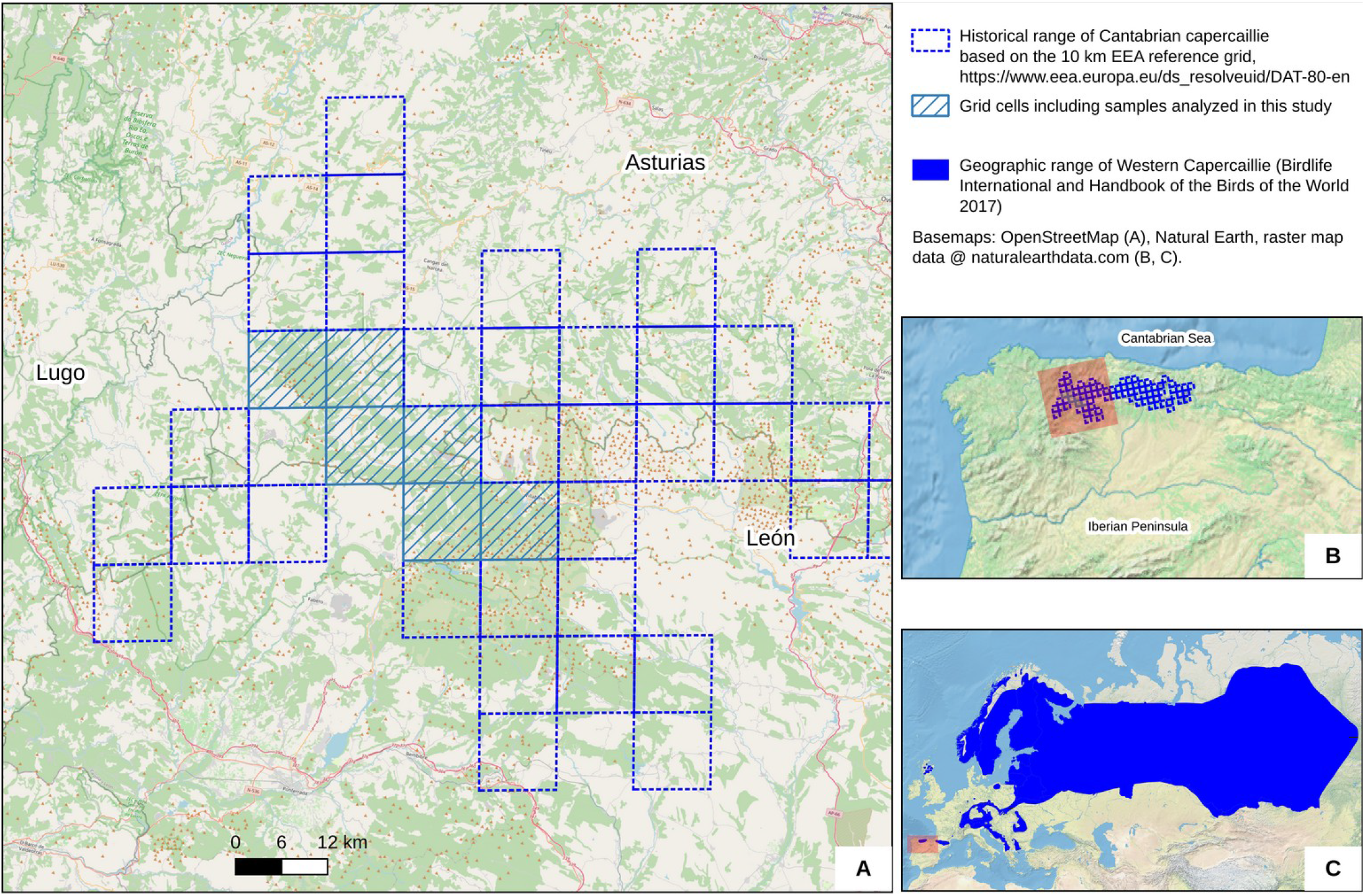
Study area.

Capercaillie gather at lekking areas in the mating season, thus making it more feasible to obtain sufficient droppings or feathers than at other times, when the birds are more dispersed and the habitat use of females and males is distinct (Bañuelos et al. 2008). We visited areas including 62 previously known capercaillie leks during the mating seasons (mid-March to early June) of 2009, 2010 and 2011 (Morán-Luis et al. 2014; Bañuelos et al. 2019 for details). Those reference locations of leks derived in fact from the period of capercaillie hunting during the mating season, which extended for much of the 20^th^ century (Rodríguez-Muñoz et al., 2015).

Of those leks, 83% showed signs of capercaillie presence at some point since year 2000; the rest had apparently been previously unused. The study area includes several largely forested areas with capercaillie presence, separated by valleys where human population and activity are mostly concentrated. Forest cover in the study area averaged 46%

Using the location of each lek from previous reports as reference, two people surveyed those forest patches for 2-3 h. Locations where we did not find signs of capercaillie presence were surveyed again 2-3 weeks later. The position of each sample was recorded with a GPS (± 5m) and was later incorporated into a GIS. To minimize redundant samples and the risk of over-sampling the same individual, we selected samples based on a minimum distance of 25 m from others with similar appearance (i.e. we weighed in freshness, size, shape and apparent content) in the case of droppings, or from the same sex in the case of feathers. Droppings were stored in tubes with silica-gel in the field, and were kept frozen at −20°C until DNA extraction. Feathers were kept dry at room temperature. We assigned samples to *display areas* instead of the historical leks used as reference because in some instances lek separation seemed an artifact, particularly considering that capercaillie display do not conform to the classical lek notion (Wegge et al. 2013).

### DNA tagging and movements

We aimed at obtaining individual genotype profiles using nine microsatellite markers (five previously developed for *Tetrao urogallus*, TUD2, TUD4, TUD5, TUT1, TUT3, Segelbacher et al. 2000; and four developed for the closely related *Tetrao tetrix*, TTD2, TTD6, BG10, BG15, Caizergues et al. 2001, Piertney and Höglund 2001), and a specific primer developed for sex assignment for Cantabrian capercaillie (Pérez et al. 2011). Details of methods of DNA extraction and amplification, molecular sexing, validation of genotype profiles and genotyping errors, were detailed in Bañuelos et al. (2019). To ensure genotype reliability, we double-checked each genotype profile. We kept only those samples for which at least six microsatellite loci amplified correctly, and which rendered a reliable and unequivocal consensus genotype (Morán-Luis et al. 2014, Bañuelos et al. 2019).

We recorded the repeated finding of capercaillie genotypes (hereafter *recaptures*) in successively processed samples. Note that the initial observation of an individual does not necessarily imply its first presence in a lek, as scats cannot accurately be dated in this context. We used individuals recaptured at least once for subsequent movement analyses. We focused particularly on the maximum planimetric distance between recaptures of each individual, both within each single mating season of 2009, 2010 and 2011, and between each mating season and the subsequent ones. We coded those recaptures as philopatric if they occurred within the same display area.

## Results

We found 127 capercaillie individuals (genotypes) in the study area: 48 females, 73 males, and 6 individuals that could not be sexed. Those capercaillies were the minimum number of individuals present in the area during the study. Individual identifications were derived from DNA extracted and amplified correctly in 422 samples, out of an initial set of 752 samples.

We found capercaillies in 35 display areas, which included 47 historical lek locations. In two of those display areas we did not find males. The maximum number of males per display area was 7, with a median of 2. The maximum number of females per display area was 4 – not coincident with the display area with most males – and the median was 1.

Seventy of the above capercaillies, 22 females (46%) and 48 (66%) males, were recaptured at least once. We recaptured 53 individuals within any single mating season, and 49 individuals between mating seasons (Fig. 2).

**Figure 2.**
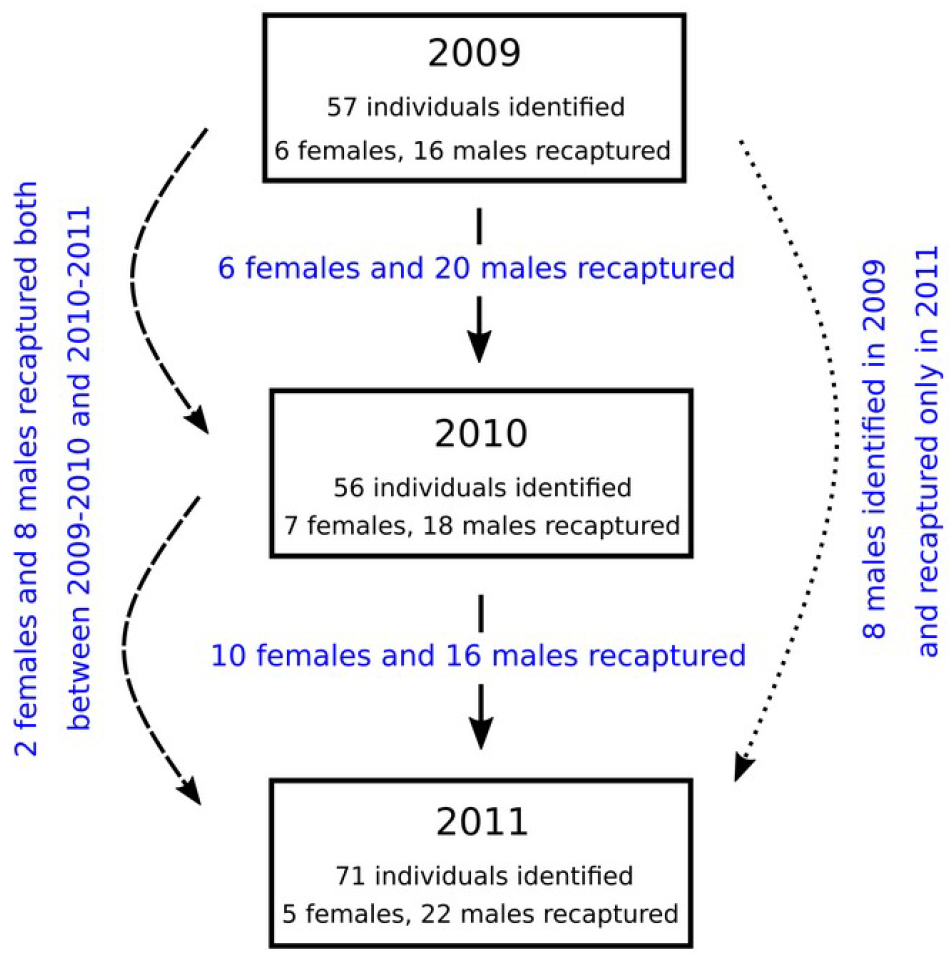
Scheme of recaptures. Scheme of capercaillie individuals identified and recaptured per season (boxes), recaptured in consecutive seasons (solid arrows), and recaptured between seasons.

Most recaptures occurred within 500 m of the initial observation, both within and between mating seasons (Fig. 3, 4). Distributions were highly skewed, including a few relatively long-distance movements (Fig. 4, 5). Table 1 shows descriptive statistics of maximum distances of recapture of each individual.

**Figure 3.**
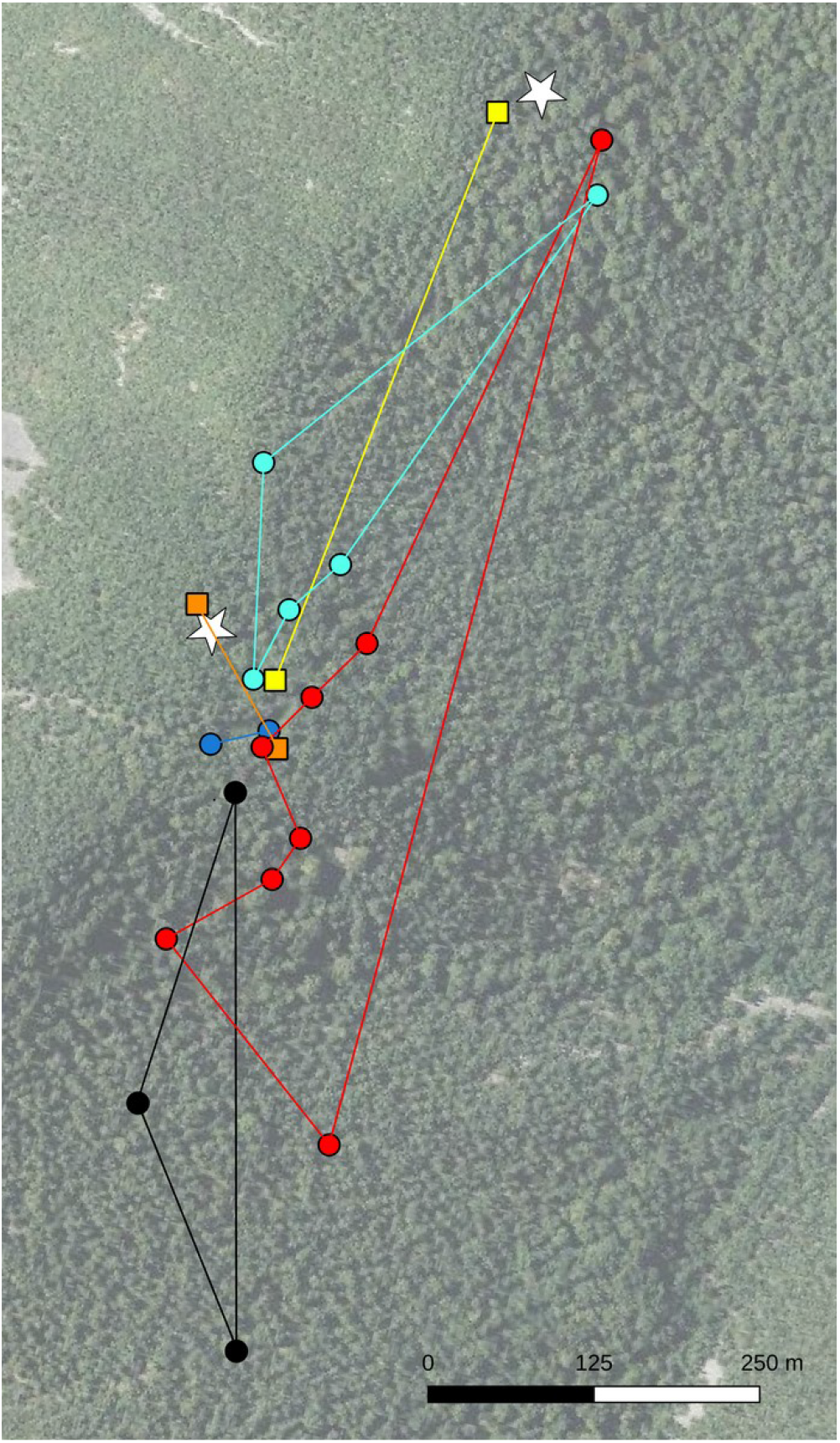
Pattern of philopatric movements within mating seasons. Example of the short movements that where the norm in a mating season. Symbols show locations of 4 capercaillie males (circles) and 2 females (squares) at a display area in 2009. Lines show idealized tracks as minimum concave polygons for each individual. White stars mark the reference location of two areas historically considered as leks. Basemap: aerial image from the Spanish Geographic Institute (https://pnoa.ign.es).

**Figure 4.**
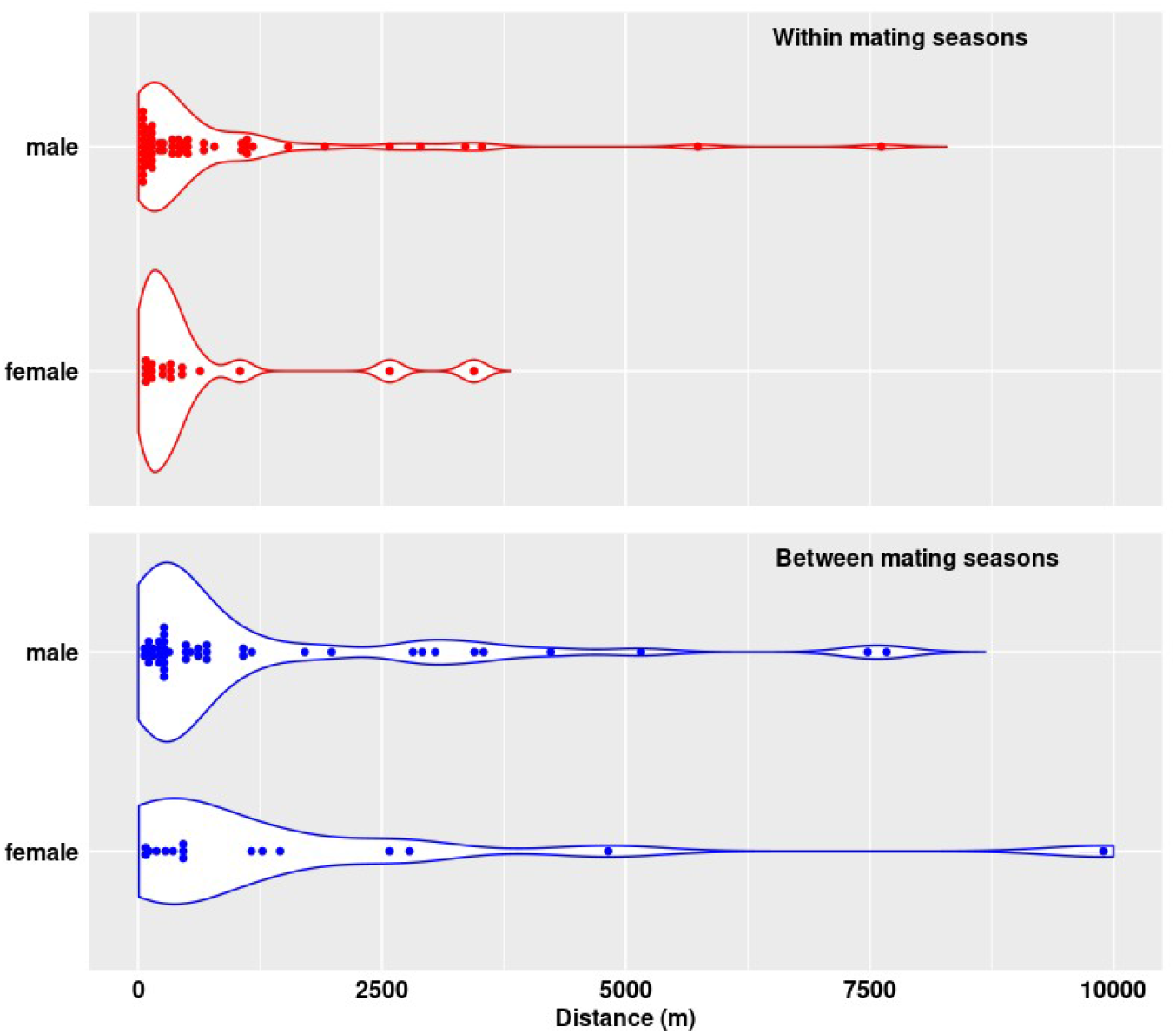
Distance of recaptures. Violin plots (data points plus a probability density) showing frequency distribution of maximum planimetric distance of recapture (m) for each capercaillie individual, both within and between mating seasons.

**Figure 5.**
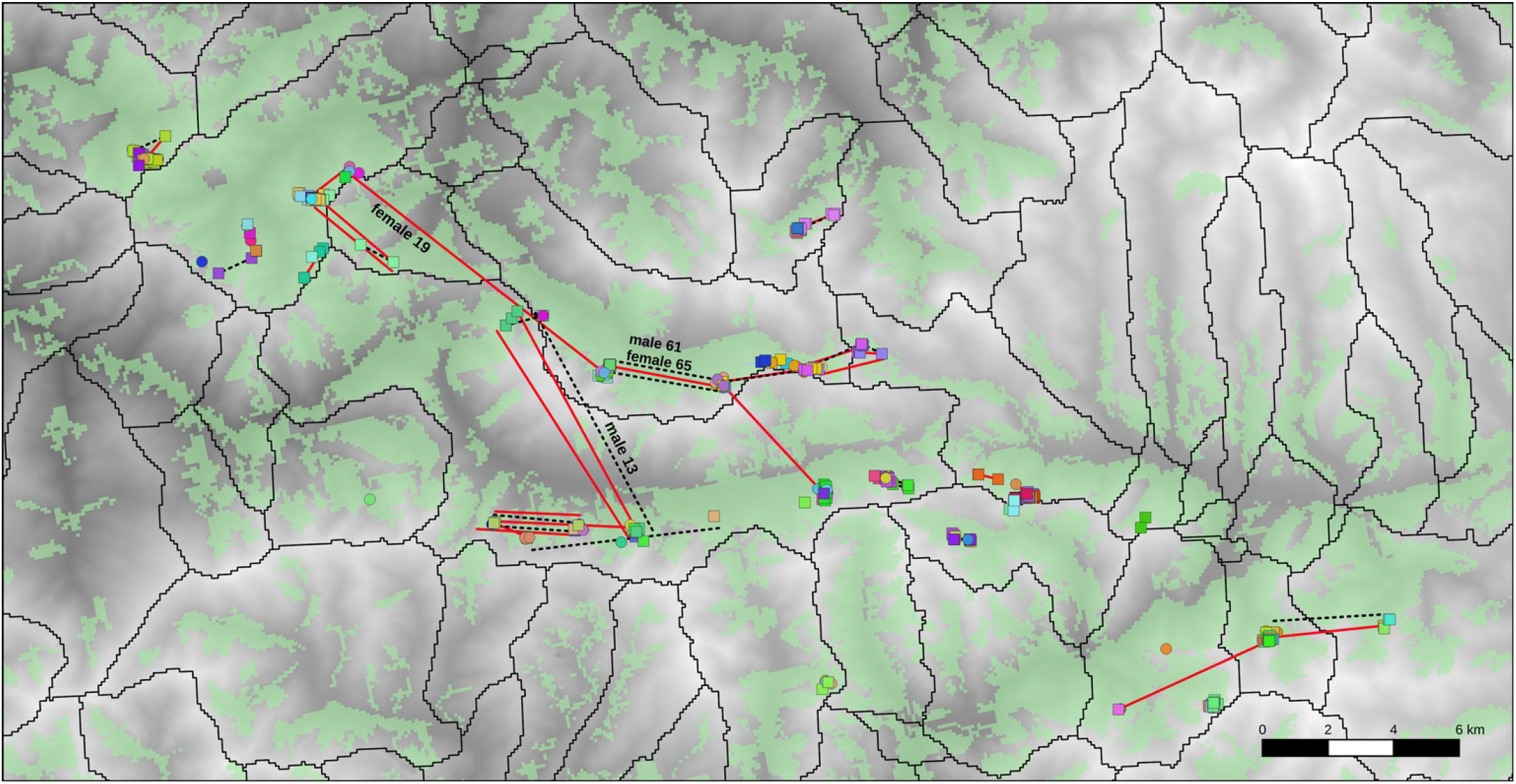
Non-philopatric recaptures and movements. Squares and circles show DNA-tagged male and female capercaillie, respectively. Lines show maximum straight line movements between recaptures of capercaillie genotypes within (solid red) and between (dashed black) mating seasons. Overlapping movements were slightly displaced for clarity. Polygons show sub-basins* over a digital elevation model of the study area where lighter hues indicate higher elevation (range 357 to 2094 m a.s.l.). *Ministerio para la T ransición Ecológica y el Reto Demográfico [https://www.miteco.gob.es/en/cartografia-y-sig/ide/descargas/agua/cuencas-y-subcuencas.aspx]

**Table 1.** Descriptive statistics of maximum distances of recapture (m) of each individual

| Period | Sex | Recaptures | Median | Median absolute deviation | Range |
| --- | --- | --- | --- | --- | --- |
| Within mating seasons | female | 18 | 297 | 252 | 63 - 3443 |
|  | male | 56 | 297 | 341 | 29 - 7618 |
| Between mating seasons | female | 16 | 483 | 599 | 59 - 9895 |
|  | male | 44 | 486 | 460 | 51 - 7673 |

There were no significant differences between females and males in maximum recapture distances within mating seasons (Wilcoxon rank sum test, W = 497, p = 0.947), or between matin seasons (Wilcoxon rank sum test, W = 375, p = 0.701). Maximum recapture distances were larger between mating seasons (Wilcoxon rank sum test, W = 2804, p = 0.01).

The farthest recapture within any mating season was that of a male that moved 7.6 km between two major valleys of the study area (Fig. 5, male 13). The farthest recapture between mating seasons was that of a female re-identified in 2011 at 9.9 km from her previous location in 2010 (Fig. 5, female 19). That female visited different display areas each spring, changing sub-basins of the study area.

In terms of philopatry, 79 % of male recaptures and 83% of female recaptures within mating seasons occurred at the same display area. 73 % of male recaptures and 56 of female recaptures between mating seasons occurred at the same display area.

About one quarter of the birds (32% of males, 23% of females) visited more than one display area at some point. There were no differences between females and males in the proportion of recaptures registered in display areas different from the initial (X^2^ = 0.08, df = 1, p = 0.78 within mating seasons; X^2^ = 1.47, df = 1, p = 0.23 between mating seasons).

Most birds that visited more than one display area during a single mating season were found at 2 different ones (3 females, 10 males); only one male was recaptured at 3 different display areas. These non-philopatric recaptures during the mating season occurred essentially between display areas located within sub-basins of the study area. We also found a coincident, non-philopatric 3.5 km movement of a female and a male in the mating season of 2010 (Fig. 5, individuals 61 and 65). A notable outlayer to the pattern just described corresponded to the male 13, mentioned above, which in the mating season of 2011 moved between two major valleys, separated by deforested terrain in southern exposure, and other human-modified terrain.

Most birds shifting display areas between mating seasons were recaptured at 2 different ones (5 females, 8 males), with just 1 female and 2 males recaptured at 3 different display areas. These recaptures also occurred mostly within major valleys, although there were three recaptures bridging sub-basins. Interestingly, two of them from 2009 to 2010, and 2010 to 2011, belonged to the same ‘travelling’ male 13 referred above (Fig. 5).

## Discussion

Surveying display areas during three consecutive breeding seasons, we tracked different capercaillie individuals in space and time. DNA-tagging recaptures provided a view of the use of space of individuals of this endangered population that we did not previously have. We found high capercaillie fidelity to display areas, but also rarer and much longer movements. Birds tended to remain in one display area in spring, although a quarter of the individuals did visit 2 or 3 display areas. In males, the latter seemed consistent with expected exploratory movements into other males’ territories (Wegge et al. 2013). Overall the prevalence of short movements in the mating season agrees with previous studies in capercaillie (e.g. Wegge and Larsen 1987, Gjerde et al. 2000), and more broadly in lekking grouse species (Cross et al. 2017). However, we expected that in this endangered population, inhabiting a fragmented forest landscape, females visited several leks in spring due to low availability of males, as suggested in the case of inter-lek movements of females in the Alps (Storch 1997). Overall, the observed pattern of movement within mating seasons implies that individuals used several historically recognised leks, in a pattern that reminded of the exploded leks notion (Wegge et al. 2013). Many of those registered leks may in fact be merged into fewer display areas, at least in terms of planning survey design and efforts.

Despite the prevalence of philopatric movements, non-philopatric recaptures over much longer distances showed that both females and males of Cantabrian capercaillie were able to move regularly through the landscape at ecological time scales. These results complement previous data on gene flow and thus longer time scales, which showed that there was no genetic subdivision among individuals living in relatively distant sub-basins of the landscape (Fameli et al. 2017). We did not find longer movements by females, despite that previous studies showed females dispersing farther, and moving more frequently between mating seasons (Segelbacher et al. 2008; Watson and Moss 2008).

Our study was focused on relatively small-scale movements. Studies that explicitly attempted to cover dispersal would in principle obtain much lower recapture rates, but would also identify longer movement events (Koenig et al. 1996; Cross et al. 2017). It is conceivable that year-round movements would be longer than those we detected in the mating season (Saniga 2006, Zizas et al. 2012). Possibly those would also be directed towards alternative habitats (e.g. Watson and Moss 2008), and could take birds outside of the study area, which is nested among historical capercaillie territories (Fig. 1). Thus, those historical territories, which today could be perceived as peripheral but which include substantial forest cover, should also be protected and monitored, as an objective for the potential recovery of the Cantabrian capercaillie population.

A related question is where were the birds that we did not recapture. Some of them were likely in the same area, and were not detected in our fieldwork, as reflected by mark-recapture models of population dynamics (Bañuelos et al. 2019). But there are other, non-exclusive possibilities. We had previously found that females showed higher turnover in the study area, and yielded less recaptures (Morán-Luis et al. 2014; Bañuelos et al. 2019). Those results could be related to lower female survival, but also to more frequent dispersal events that remained undetected in our sampling schemes. It is possible that birds kept dispersing outside of the study area, which we selected for its consistent capercaillie presence in the recent decades. Data from Russian capercaillie populations suggested that younger birds switched an area of primary habitat for lower quality, logged neighbouring patches, due to competition with dominant adults, and some returned later as adults (Borchtchevski 1993). An equivalent pattern of sources and sinks in capercaillie local populations has been described in the Bavarian Alps (Segelbacher et al. 2003). We speculate that a similar process could be taking place in our study area, which includes higher forest cover and quality than the adjacent landscape (Quevedo et al. 2006). However, evaluating that idea would require regular, formal surveying efforts, and we are not aware of any. Capercaillie management in the Cantabrian Mountains has paid little if any attention to forest patches outside the historically known display areas. Moreover, as the presence of capercaillie in spring declined in historically known territories, attention has been placed on a subsequently smaller fraction of those, without monitoring formerly used forests, or potential new locations.

### Ups and downs of studying movements with DNA tagging

Comparing different aspects of tracking animals based on DNA-tagging vs. direct tracking is not straightforward, yet we saw both pros and cons of DNA-tagging: We identified 127 individuals, and registered movements for 70 of them. Such indirect tracking thus provided a much larger sample size of *tagged* individuals distances than could possibly be obtained by a standard research team working with endangered birds, and under standard Spanish research funding. In addition, DNA-tagging based on scats is a non invasive technique, particularly suited to individuals of endangered populations, for which handling stress would be a serious problem (Gibson et al. 2013; Blomberg et al. 2018). Results may also be less influenced by behavioral bias, like avoidance of the observer, or response to handling and initial mark (Miller et al. 2005; Biro and Dingemanse 2009; Garamszegi et al. 2009).

On the other hand, what we gained in number of individuals identified, we certainly missed in the number of data points per individual. Present-day GPS tracking technology could provide a much larger datasets, thus allowing detailed inferences (e.g. de Gabriel Hernando et al. 2020). Another shortcoming is that using DNA-tagging based on shed tissues or scats it is not possible detailing the sequence of the initial and subsequent “captures”; thus it is not possible tracing precise movement paths. In addition, we lack direct information on the natural history of the “tagged” birds. We lacked for instance important individual information like age and social status; aspects that would have been useful to interpret whether the registered movements were dispersive or exploratory. Or to check whether results were biased to adult males. Such bias is certainly possible, because the presence of subordinate males may be more difficult to detect in spring (Wegge and Larsen 1987), even though we had it in mind, and included as much surveyed terrain as possible in the periphery of reference lek locations.

## Acknowledgments

The study was funded by grants IB08-158 (FICYT, Asturian Government) to MJ Bañuelos and CGL2010-15990 (MICINN, Spanish Government) to MJ Bañuelos and M Quevedo. We thank Alberto Fernández-Gil, Bea Blanco Fontao, and Rolando Rodríguez Muñoz for their help in designing and conducting field surveys. Eduardo González, Víctor Rodríguez, and Damián Ramos helped with the fieldwork, and Jose Carral helped with field logistics. Alberto Fameli greatly helped fine-tuning lab methods, and Leticia Viesca helped analysing individual genotypes. We acknowledge the permits granted by the environmental authorities of Asturias and León, required to survey capercaillie mating areas.

## References

Baguette M, Blanchet S, Legrand D, Stevens VM, Turlure C. 2013. Individual dispersal, landscape connectivity and ecological networks. Biological Reviews 88(2):310–326. doi:10.1111/brv.12000.

Bañuelos MJ, Blanco-Fontao B, Fameli A, Fernández-Gil A, Mirol P, Morán-Luis M, Rodríguez-Muñoz R, Quevedo M. 2019. Population dynamics of an endangered forest bird using mark-recapture models based on DNA-tagging. Conservation Genetics 20(6):1251–1263. doi:10.1007/s10592-019-01208-x.

Bañuelos MJ, Quevedo M, Obeso JR. 2008. Habitat partitioning in endangered Cantabrian capercaillie *Tetrao urogallus cantabricus*. Journal of Ornithology 149:245–252. doi:10.1007/s10336-007-0267-5.

Biro PA, Dingemanse NJ. 2009. Sampling bias resulting from animal personality. Trends in Ecology & Evolution 24(2):66–67. doi:10.1016/j.tree.2008.11.001.

Blanco-Fontao B, Fernández-Gil A, Obeso J, Quevedo M. 2010. Diet and habitat selection in Cantabrian Capercaillie (*Tetrao urogallus cantabricus*): ecological differentiation of a rear-edge population. Journal of Ornithology 151(2):269–277. doi:10.1007/s10336-009-0452-9.

Blomberg E, Davis S, Mangelinckx J, Sullivan K. 2018. Detecting capture-related mortality in radio-marked birds following release. Avian Conservation and Ecology 13(1). doi:10.5751/ACE-01147-130105.

Bonte D, Van Dyck H, Bullock JM, Coulon A, Delgado M, Gibbs M, Lehouck V, Matthysen E, Mustin K, Saastamoinen M. 2012. Costs of dispersal. Biological Reviews 87(2):290–312. doi:10.1111/j.1469-185X.2011.00201.x

Borchtchevski VG. 1993. Population biology of the capercaillie. Principles of the structural organization [in Russian; tables and plots in English]. Moscow: Central Laboratory of the Management of Hunting and Nature Reserves.

Börger L, Dalziel BD, Fryxell JM. 2008. Are there general mechanisms of animal home range behaviour? A review and prospects for future research. Ecology Letters 11(6):637–650. doi:10.1111/j.1461-0248.2008.01182.x.

Caizergues A, Dubois S, Loiseau A, Mondor G, Rasplus J-Y. 2001. Isolation and characterization of microsatellite loci in black grouse (*Tetrao tetrix*). Molecular Ecology Notes 1(1-2):36–38. doi:10.1046/j.1471-8278.2000.00015.x.

Caplat P, Edelaar P, Dudaniec RY, Green AJ, Okamura B, Cote J, Ekroos J, Jonsson PR, Löndahl J, Tesson SV, et al. 2016. Looking beyond the mountain: dispersal barriers in a changing world. Frontiers in Ecology and the Environment 14(5):261–268. doi:10.1002/fee.1280.

Cervellini M, Zannini P, Musciano MD, Fattorini S, Jiménez-Alfaro B, Rocchini D, Field R, Vetaas OR, Irl SDH, Beierkuhnlein C, et al. 2020. A grid-based map for the Biogeographical Regions of Europe. Biodiversity Data Journal 8:e53720. doi:10.3897/BDJ.8.e53720.

Channel R, Lomolino MV. 2000. Dynamic biogeography and conservation of endangered species. Nature 403 (6765):84–86. doi:10.1038/47487.

Cross TB, Naugle DE, Carlson JC, Schwartz MK. 2017. Genetic recapture identifies long-distance breeding dispersal in Greater Sage-Grouse (*Centrocercus urophasianus*). The Condor 119(1):155–166. doi:10.1650/CONDOR-16-178.1.

de Gabriel Hernando M, Karamanlidis AA, Grivas K, Krambokoukis L, Papakostas G, Beecham J. 2020. Reduced movement of wildlife in Mediterranean landscapes: a case study of brown bears in Greece. Journal of Zoology 311(2):126–136. doi:10.1111/jzo.12768

Fameli A, Morán-Luis M, Rodríguez-Muñoz R, Bañuelos MJ, Quevedo M, Mirol P. 2017. Conservation in the southern edge of *Tetrao urogallus* distribution: Gene flow despite fragmentation in the stronghold of the Cantabrian capercaillie. European Journal of Wildlife Research 63(3). doi:10.1007/s10344-017-1110-9.

Garamszegi LZ, Eens M, Török J. 2009. Behavioural syndromes and trappability in free-living collared flycatchers, *Ficedula albicollis*. Animal Behaviour 77(4):803–812. doi:10.1016/j.anbehav.2008.12.012.

Gibson D, Blomberg EJ, Patricelli GL, Krakauer AH, Atamian MT, Sedinger JS. 2013. Effects of radio collars on survival and lekking behavior of male Greater Sage-Srouse. The Condor. 115(4):769–776. doi:10.1525/cond.2013.120176.

Gjerde I, Wegge P, Rolstad J. 2000. Lost hotspots and passive female preference: the dynamic process of lek formation in capercaillie *Tetrao urogallus*. Wildlife Biology 6(4):291–298. doi:10.2981/wlb.2000.029.

Koenig WD, Van Vuren D, Hooge PN. 1996. Detectability, philopatry, and the distribution of dispersal distances in vertebrates. Trends in Ecology & Evolution 11(12):514–517. doi:10.1016/S0169-5347(96)20074-6.

Lechner AM, Doerr V, Harris RMB, Doerr E, Lefroy EC. 2015. A framework for incorporating fine-scale dispersal behaviour into biodiversity conservation planning. Landscape and Urban Planning 141:11–23. doi:10.1016/j.landurbplan.2015.04.008.

Lens L, Van Dongen S, Norris K, Githiru M, Matthysen E. 2002. Avian persistence in fragmented rainforest. Science 298(5596):1236–1238. doi:10.1126/science.1075664.

Miller CR, Joyce P, Waits LP. 2005. A new method for estimating the size of small populations from genetic mark–recapture data. Molecular Ecology 14(7):1991–2005. doi:10.1111/j.1365-294X.2005.02577.x.

Ministerio para la Transición Ecológica. 2018. Declaración de situación crítica de Cistus heterophyllus subsp. carthaginensis, Lanius minor, Margaritifera auricularia, Marmaronetta angustirostris, Mustela lutreola, Pinna nobilis y Tetrao urogallus cantabricus en España. https://www.boe.es/eli/es/o/2018/09/28/tec1078.

Morán-Luis M, Fameli A, Blanco-Fontao B, Fernández-Gil A, Rodríguez-Muñoz R, Quevedo M, Mirol P, Bañuelos MJ. 2014. Demographic status and genetic tagging of endangered capercaillie in NW Spain. PLOS ONE 9(6):e99799. doi:10.1371/journal.pone.0099799.

Olson DM, Dinerstein E, Wikramanayake ED, Burgess ND, Powell GVN, Underwood EC, D’amico JA, Itoua I, Strand HE, Morrison JC, et al. 2001. Terrestrial ecoregions of the world: a new map of life on Earth. BioScience 51(11):933. doi:10.1641/0006-3568(2001)051[0933:TEOTWA]2.0.CO;2.

Palsbøll PJ. 1999. Genetic tagging: contemporary molecular ecology. Biological Journal of the Linnean Society 68(1–2):3–22. doi:10.1111/j.1095-8312.1999.tb01155.x.

Palsbøll PJ, Allen J, Bérubé M, Clapham PJ, Feddersen TP, Hammond PS, Hudson RR, Jørgensen H, Katona S, Larsen AH, et al. 1997. Genetic tagging of humpback whales. Nature 388(6644):767–769. doi:10.1038/42005.

Pérez T, Vázquez JF, Quirós F, Domínguez A. 2011. Improving non-invasive genotyping in capercaillie (*Tetrao urogallus):* redesigning sexing and microsatellite primers to increase efficiency on faeces samples. Conservation Genetics Resources 3(3):483–487. doi:10.1007/s12686-011-9385-8.

Piertney SB, Höglund J. 2001. Polymorphic microsatellite DNA markers in black grouse (*Tetrao tetrix*). Molecular Ecology Notes 1(4):303–304. doi:10.1046/j.1471-8278.2001.00118.x.

Pollo C, Robles L, Seijas JM, García-Miranda A, Otero R. 2005. Trends in the abundance of Cantabrian Capercaillie *Tetrao urogallus cantabricus* at leks on the southern slope of the Cantabrian Mountains, north-west Spain. Bird Conservation International 15(04):397–409. doi:10.1017/S0959270905000626.

Quevedo M, Bañuelos MJ, Obeso JR. 2006. The decline of Cantabrian capercaillie: How much does habitat configuration matter? Biological Conservation 127(2):190–200. doi:10.1016/j.biocon.2005.07.019.

Ricketts TH. 2001. The matrix matters: effective isolation in fragmented landscapes. The American Naturalist 158(1):87–99. doi:10.1086/320863.

Rodríguez-Muñoz R, del Valle CR, Bañuelos MJ, Mirol P. 2015. Revealing the consequences of male-biased trophy hunting on the maintenance of genetic variation. Conservation Genetics. 16:1375–1394. doi:10.1007/s10592-015-0747-8.

Sandel B, Svenning J-C. 2013. Human impacts drive a global topographic signature in tree cover. Nature Communications 4. doi:10.1038/ncomms3474.

Saniga M. 2006. Home range sizes and roosting places in capercaillie (*Tetrao urogallus* L.) cocks living solitary in the West Carpathians. Folia Oecologica 33(2): 121–128.

Segelbacher G, Manel S, Tomiuk J. 2008. Temporal and spatial analyses disclose consequences of habitat fragmentation on the genetic diversity in capercaillie (*Tetrao urogallus*). Molecular Ecology 17:2356–2367. doi:10.1111/j.1365-294X.2008.03767.x.

Segelbacher G, Paxton RJ, Steinbrück G, Trontelj P, Storch I. 2000. Characterization of microsatellites in capercaillie *Tetrao urogallus* (Aves). Molecular Ecology 9:1934–1936. doi:10.1046/j.1365-294x.2000.0090111934.x.

Segelbacher G, Storch I, Tomiuk J. 2003. Genetic evidence of capercaillie *Tetrao urogallus* dispersal sources and sinks in the Alps. Wildlife Biology 9(4):267–273. doi:10.2981/wlb.2003.014.

Southwell DM, Lechner AM, Coates T, Wintle BA. 2008. The sensitivity of population viability analysis to uncertainty about habitat requirements: implications for the management of the endangered southern brown bandicoot. Conservation Biology 22(4):1045–1054. doi:10.1111/j.1523-1739.2008.00934.x.

Storch I. 1997. Male territoriality, female range use, and spatial organisation of capercaillie *Tetrao urogallus* leks. Wildlife Biology 3(3-4):149–161. doi:10.2981/wlb.1997.019.

Storch I, Bañuelos MJ, Fernández-Gil A, Obeso JR, Quevedo M, Rodríguez-Muñoz R. 2006. Subspecies Cantabrian capercaillie *Tetrao urogallus cantabricus* endangered according to IUCN criteria. Journal of Ornithology 147(4):653–655. doi:10.1007/s10336-006-0101-5.

Watson A, Moss R. 2008. Grouse: The Natural History of British and Irish Species. 1st edition. London: Collins.

Watson JEM, Venter O, Lee J, Jones KR, Robinson JG, Possingham HP, Allan JR. 2018. Protect the last of the wild. Nature 563(7729): 27. doi:10.1038/d41586-018-07183-6.

Wegge P, Larsen B. 1987. Spacing of adult and subadult male common Capercaillie during the breeding season. The Auk 104:481–490. doi:10.2307/4087547.

Wegge P, Rolstad J, Storaunet KO. 2013. On the spatial relationship of males on ‘exploded’ leks: the case of Capercaillie grouse *Tetrao urogallus* examined by GPS satellite telemetry. Ornis Fennica 90(4):222–235.

Yoder JM, Marschall EA, Swanson DA. 2004. The cost of dispersal: predation as a function of movement and site familiarity in ruffed grouse. Behavioral Ecology 15(3):469–476. doi:10.1093/beheco/arh037.

Zizas R, Shamovich D, Kurlavičius P, Belova O, Brazaitis G. 2012. Radio-tracking of Capercaillie (*Tetrao urogallus* L.) in North Belarus. Baltic Forestry 18(2):270–277.

